# Impact of growth-promoting alternatives on weight gain and gut microbial diversity and activity in piglets

**DOI:** 10.1101/2022.12.12.520170

**Authors:** Jason Palanee, Nathalie Gagnon, Karoline Lauzon, Steve Méthot, Mylène Blais, Guylaine Talbot, Frédéric Guay, Martin Lessard, Étienne Yergeau, Dominic Poulin-Laprade

**Author notes:** Correspondence: Dominic Poulin-Laprade, 2000 College Street, Sherbrooke, QC J1M 1Z3, Canada.

## Abstract

In swine husbandry, weaning is a critical event for piglets which causes environmental, nutritional, and psychological stresses, with consequences such as intestinal dysbiosis. To counteract this issue, producers resorted to the use of in-feed antimicrobials to prevent post-weaning diarrhea and to promote growth for increased animal performance. However, the use of antibiotic for growth promotion was banned in many countries. In-feed supplements have great potential as alternative strategies. This study evaluated the effect on gut microbial activity, microbiome, and animal performance of combinations of peri-weaning feeding strategies such as bovine colostrum, medium-chain fatty acids and yeast extract. We quantified weight gain, intestinal pH, volatile fatty acids, and characterized the gut microbiota on ileum, cecum, and colon digestates. Overall, the feed supplements had limited impact on weight gain and volatile fatty acids production. However, the combined treatments have demonstrated a modulatory effect on gut microbiota which supports a potential role as an alternative to growth-promoting antibiotic in the swine industry.

## Introduction

In the swine industry, weaning has many detrimental consequences on piglets’ health caused by abrupt dietary and environmental changes (Campbell et al., 2013;Gresse et al., 2017). At the time of weaning, the piglets will transition from lactation to a solid vegetable-based diet which promotes weight gain (Heo et al., 2013). In addition to this feeding transition, the piglets are separated from the sow and are introduced into a new pen where they must establish the hierarchical order again (Barba-Vidal et al., 2018). Weaning is intensely stressful for the piglets that still have an immature intestinal immune system and microbiota, resulting in numerous physiological, immunological, and microbiological changes causing reduced animal performance, increased susceptibility to enteric infection and in some cases, mortality (Campbell et al., 2013;Heo et al., 2013;Gresse et al., 2017;Barba-Vidal et al., 2018).

The conditions surrounding weaning often involve the development of intestinal dysbiosis which weakens the barrier role of the intestinal mucosa, making it more permeable to allergens and bacteria such as opportunistic pathogens resulting in enteric infections (Wijtten et al., 2011;Rhouma et al., 2017). These enteric infections are the cause of post-weaning diarrhea often associated with the proliferation of enterotoxigenic *Escherichia coli* strains in the gastrointestinal tract (Rhouma et al., 2017). To reduce the effects of weaning, cases of infections, and important economic losses, producers resorted to the use of in-feed antibiotics (Gresse et al., 2017;Barba-Vidal et al., 2018). In addition to the prophylaxis and therapeutic aspect, antibiotics were commonly used in the Canadian and worldwide swine industry as growth promoters (Campbell et al., 2013;Holman and Chénier, 2015;Gresse et al., 2017). Canada and many other countries have banned the use of medically important antibiotic for growth promotion, making the search for alternative growth promoters a crucial need (Barba-Vidal et al., 2018).

Currently many research teams are studying the potential of feed supplements as alternatives to the use of in-feed antibiotics (Verstegen and Williams, 2002). Different feed supplements promise to control the presence of pathogens while maintaining good intestinal health and promoting animal performance (Langlais, 2019). Among the different types of known feed supplements, colostrum, medium-chain fatty acids, yeast appeared to be good candidates because of their very interesting complementary functional properties. Indeed, these compounds have together the potential to promote antimicrobial and immunoregulatory activities, as well as growth performance. Bovine colostrum contains bioactive molecules with immunoregulatory and antimicrobial properties that are essential for the development of the immune system (Cross and Gill, 2000;Schlimme et al., 2000;Sugiharto et al., 2015;Lo Verso et al., 2020a;Yan et al., 2021) and modulating the inflammatory response to pathogens (Blais et al., 2015). The interest surrounding the supplementation of yeast extract is driven by its content in active components such as enzyme, nucleic acids, and cell wall products such as β-glucan and mannans. These polysaccharides have shown complementary prebiotic, antimicrobial and immunological effects (Volman et al., 2008;Lo Verso et al., 2020b). Finally, medium chain fatty acids and naked oats are able to increase appetite resulting in a better animal performance (Lauridsen, 2020;Świątkiewicz et al., 2020). In addition, medium-chain fatty acids are an excellent source of energy, have antimicrobial properties (Skřivanová et al., 2009) and allow for proper intestinal development (Hanczakowska et al., 2013;Świątkiewicz et al., 2020).

However, it is not clear how these supplements compare to antibiotics in their capacity to modulate the gut microbiota and if this affects important gut microbiota functions, such as the production of volatile fatty acids (VFA). When shifting from a liquid to a solid grain-based diet at weaning, the presence of VFA-producing bacteria is crucial to metabolize the non-digestible carbohydrates and maximize piglets’ weight gain (Zhou et al., 2019;Jiao et al., 2020;Deleu et al., 2021). The presence of VFA-producing bacteria such as *Bacteroidetes, Eubacterium* and *Coprococcus*, among others, allows the production of acetate, propionate, and butyrate, which are the three main volatile fatty acids produced by the gut microbiota (Deleu et al., 2021). The production of acetate is generally well spread among the various bacterial classes, unlike butyrate and propionate that are rather specific to the substrates (Deleu et al., 2021). There are two possible metabolic pathways to produce propionate: the succinate pathway used by the *Bacteroidetes* and some *Firmicutes*, and the propanediol pathway used by the *Lachnospiraceae*. Butyrate is produced by the butyrate kinase (*Coprococcus* genus), or by the butyryl CoA:acetate CoA transferase, (e.g. *Eubacterium*) (Deleu et al., 2021). VFAs have several beneficial effects on the metabolism (Jiao et al., 2020). They are the most important source of energy for colonocytes, allowing their growth and proliferation and thus reducing the harmful effects of weaning on colon physiology (Zhou et al., 2019).

Our hypothesis was that by their synergic effect on the piglet gut microbiota and intestinal health, a naked oat base feed supplemented with combinations of bovine colostrum, yeast extract and medium chain fatty acids will result in increased intestinal VFA concentrations, and consequently increased weight gains, to levels similar to piglets eating a chlortetracycline feed. Thus, the overall objective of this study was to evaluate the potential of the feed supplements on the development of the intestinal microbiota, its apparent VFA production and on the weight gain of post-weaning low and high birth weight piglets.

## Materials and Methods

### Animals and experimental design

Experimental procedures followed the “Canadian Council of Animal Care” guide for the care and use of farm animals in research, teaching, and testing (2^nd^ edition), and were approved by the Institutional Animal Care Committee of the Sherbrooke Research and Development Centre of Agriculture and Agri-Food Canada. All animals were cared for and slaughtered in accordance with the National Farm Animal Care Council’s code of practice for the care and handling of pigs (2014 edition). Seventy sows were used to conduct this project, for which various samples were collected between October 2018 and February 2020. At piglet’s birth, sows and their litter (12 piglets per litter) were assigned to one of five experimental treatments in a randomized complete block design. In summary, there were 13 blocks of five litters each with one litter per treatment per block. Dietary treatments, from day 7 until day 35 of age, were: 1) a base feed of 35% naked oats (CTRL); 2) CTRL feed supplemented with 5% bovine colostrum (BC); 3) CTRL feed supplemented with 0.08% medium chain fatty acid and 0.08% yeast extract (MCFA-YE); 4) CTRL feed supplemented with a combination of BC and MCFA-YE (BC-MCFA-YE); and 5) CTRL feed supplemented with 0.10% chlortetracycline (CTC). The dietary treatments were administered first through creep feeding starting at day 7 of age. Then, the same feed was served to the animals for the first two weeks after weaning at 22 days of age. On the day 35 of age, all piglets were fed with the same piglet growth diet until the end of the nursery phase. Among piglets of each litter, two low birth weights (LW) ranging between 1.0-1.25 kg and two high birth weights (HW) ranging between 1.5-1.7 kg were selected. The weights of piglets were recorded at birth and on days 1, 7, 14, 22, 26, 35, and 42 of age.

### Euthanasia of piglets

Euthanasia was performed at weaning (day 22 of age) or 4 days post-weaning (day 26) by exsanguination after the piglets were tranquilized with an injection of Stresnil (2.5 mg/kg) and stunned with a matador-type slaughter gun. Once euthanized, the contents of the ileum, cecum, and colon were collected and kept on ice until being stored at -80°C. The pH of the cecum and colon digestates were measured.

### Analysis of volatile fatty acids (VFA)

The analysis of volatile fatty acids was performed using Perkin-Elmer gas chromatograph, model Clarus 580 (Perkin-elmer Corporation, Shelton, CT, USA) coupled to a high-resolution DB-FFAP column connected to a flame ionization detector.

### DNA extraction

DNA extraction was performed using the QIAamp® DNA Stool Mini Kit (QIAGEN, Hilden, Germany) according to the manufacturer’s instruction. The DNA was extracted from digestates using a bead beating method (Zaheer et al., 2018) with minor modifications: frozen fecal sample was diluted with 0.9% saline (1:2) before processing, and the homogenization step was done in a Bead Ruptor at 4,000 rpm during 3 minutes. For all samples, the DNA integrity was verified by electrophoresis in 1% agarose gel. and quantified using the NanoDrop® apparatus and the Quant-iT PicoGreen® kit (Invitrogen)

### DNA sequencing

To characterize the gut microbiota of the piglets, the DNA samples were transferred to the Centre d’expertise et de service Genome Quebec (Montreal, Canada), where amplicon library preparation and DNA sequencing by Illumina MiSeq was performed. During sequencing, a cycle 500 kit with 250 base pair in both directions was done. No controls were performed during sequencing; however, no template controls were performed during PCR analysis, and these showed no amplification. The bacterial V4 hypervariable region of the 16S rRNA gene was amplified using the primer set 515F (5’-GTGCCAGCMGCCGCGGTAA-3’) and 806R (5’-GGACTACHVGGGTWTCTAAT-3’) (Wasimuddin et al., 2020;Correa-García et al., 2021). Based on different studies, this prokaryotic primer pair produced the highest estimations of 25 species richness and diversity in all sample categories, as well as the most diversity coverage of reference databases in in silico primer analysis (Wasimuddin et al., 2020). Raw sequence reads were deposited in the NCBI under accession PRJNA00000 (to be submitted).

### Bioinformatic analyses

Bioinformatic analyses were performed in the R environment (R_Core_Team, 2021). The demultiplexed reads were analyzed using the DADA2 bioinformatic pipeline version 1.16 in which reads went through a quality profile inspection and were then trimmed based on the error rates calculated by the parametric error model (Callahan et al., 2016;Barnes et al., 2020;Wasimuddin et al., 2020). After applying the core sample inference algorithm to the filtered and trimmed sequence data, the paired reads were merged to be able to build the amplicon sequence variants (ASV) table. The chimeras were removed and the sequence variants were assigned to the accurate taxonomy using the naive Bayesian classifier method against the Silva v138.1 nonredundant database (Quast et al., 2012;Callahan et al., 2016;Barnes et al., 2020).

### Statistical analyses

Statistical analysis on weight gain, intestinal pH, and volatile fatty acid concentration was performed using SAS (SAS Statistical Analysis System, Release 9.4, 2002-2012. SAS Institute Inc., Cary, NC). A repeated measures analysis of variance was conducted on weight gains with a randomized complete block design (N=410) by using the mean values of piglet weight per litter. Since there was a significant interaction between the treatments and the age, the differences between the treatments were considered as dissimilar at each age. An analysis at each age was therefore necessary to properly interpret this interaction.

As for weight gains, the effect of feeding treatment on intestinal pH values was tested on the average of two piglets of the same litter. Piglets selected for slaughter were selected based on birth weight (BW), so the BW effect is a subplot factor on the litter. Age at slaughter is a factor of the same type as slaughter weight. BW and age are therefore within-litter factors. The sampling site, on the other hand, corresponds to a repeated measurement on the sample unit that is the piglet. The presence of several interactions in these results requires an analysis for each site separately. Further analyses with a simpler model (split-plot) shows differences between treatments for almost every comparisons. The lack of interaction indicates that the differences between the treatments are similar for low-birth-weight piglets (LW) and high birth weight piglets (HW). Volatile fatty acids (VFA) measurements were presented in mg/L. For consistency in multivariate analysis (PCA) and univariate analyses of variance, these data have been transformed to percentages of the total (sum of VFAs). The analysis of variance of the full model on the total VFAs showed that separate analyses for each site were required. In addition, an analysis of variance for each site and each age separately was done, as well as an analysis of variance on each VFA.

R version 4.1.0 was used for statistical analysis of the amplicon sequencing data. We calculated the microbial alpha diversity (Shannon index and the number of observed species) using the otuSummary package (Yang, 2018). We evaluated the normality of our data with a Shapiro-Wilk test, and since the distribution did attain normality, we performed a univariate analysis of variance using ANOVA followed by Tukey’s honestly significant difference post hoc tests. PERMANOVA analyses tested for the effect of the treatments and their interaction on the microbial communities with 9999 permutations using the adonis function of the vegan package (Oksanen, 2011). Non-metric multidimensional scaling (NMDS) based on Bray-Curtis dissimilarity index was used to visualize variations in community composition between treatments using the vegan package. Finally, the ggplot2 package was used to produce figures (Wickham et al., 2019).

## Results

### Animal performance

The piglets were weighed for a total of seven times at different days of age, i.e. at days 1, 7, 14, 22, 26, 35 and 42. All piglets followed a similar growth curve (Fig. 1), with the exception of the piglets of the antibiotic control group (CTC), which, at days 35 and 42 of age, were heavier than the piglets of the other treatment groups (D35: P = 0.0002 and D42: P = 0.0018).

**Figure 1.**
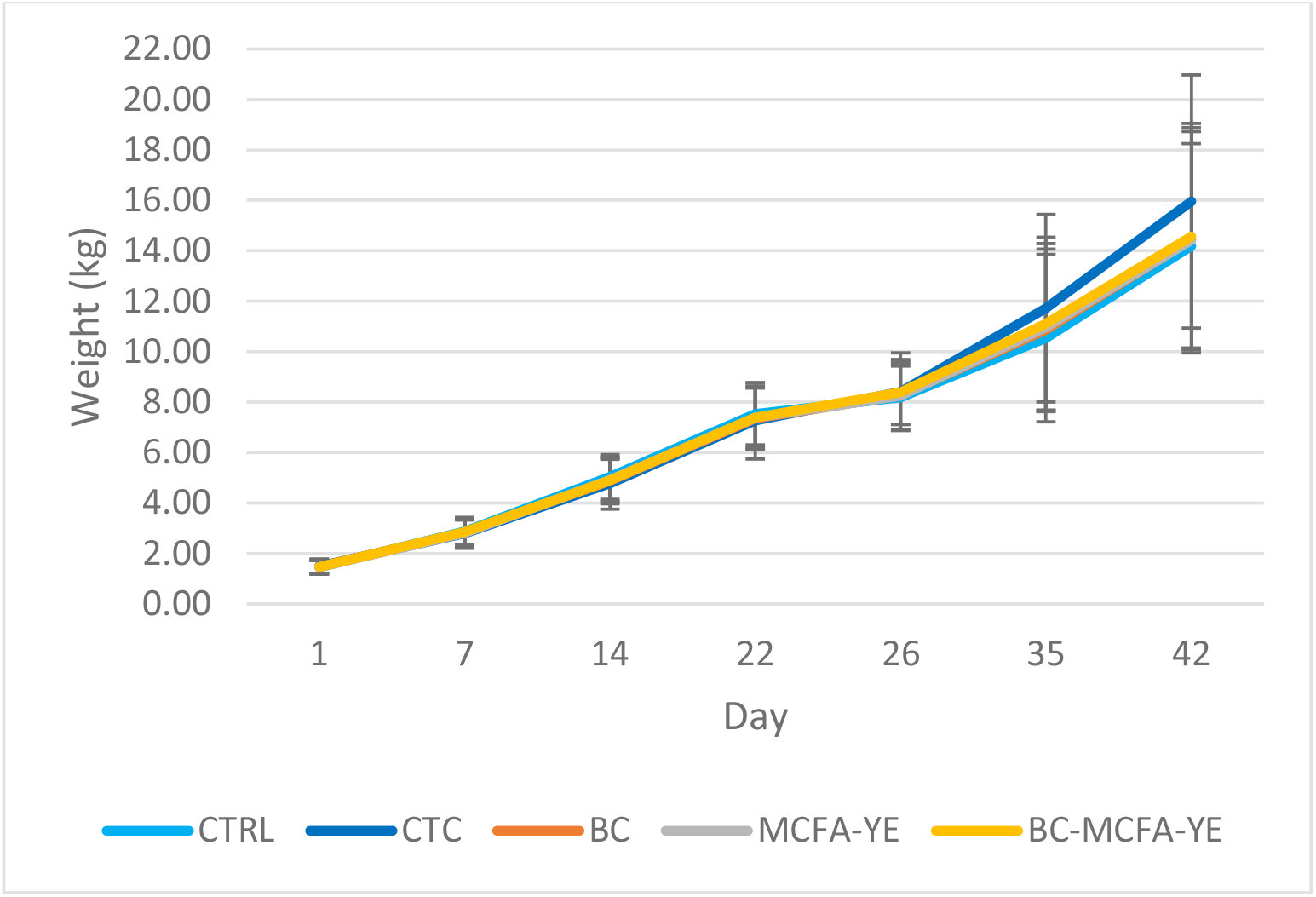
Animal performance of piglets under feed treatments from day 1 to day 42 of age.

### Intestinal pH

The pH of the cecal digestates were similar for all animals (Fig. 2a). In contrast, the pH of the colonic digestates were significantly lower in the piglets receiving the BC-MCFA-YE treatment as compared to the CTC (P < 0.05) and MCFA-YE (P < 0.05) piglets. The piglet’s birth weight had no significant effect on the pH of the intestinal content of the cecum or the colon.

**Figure 2.**
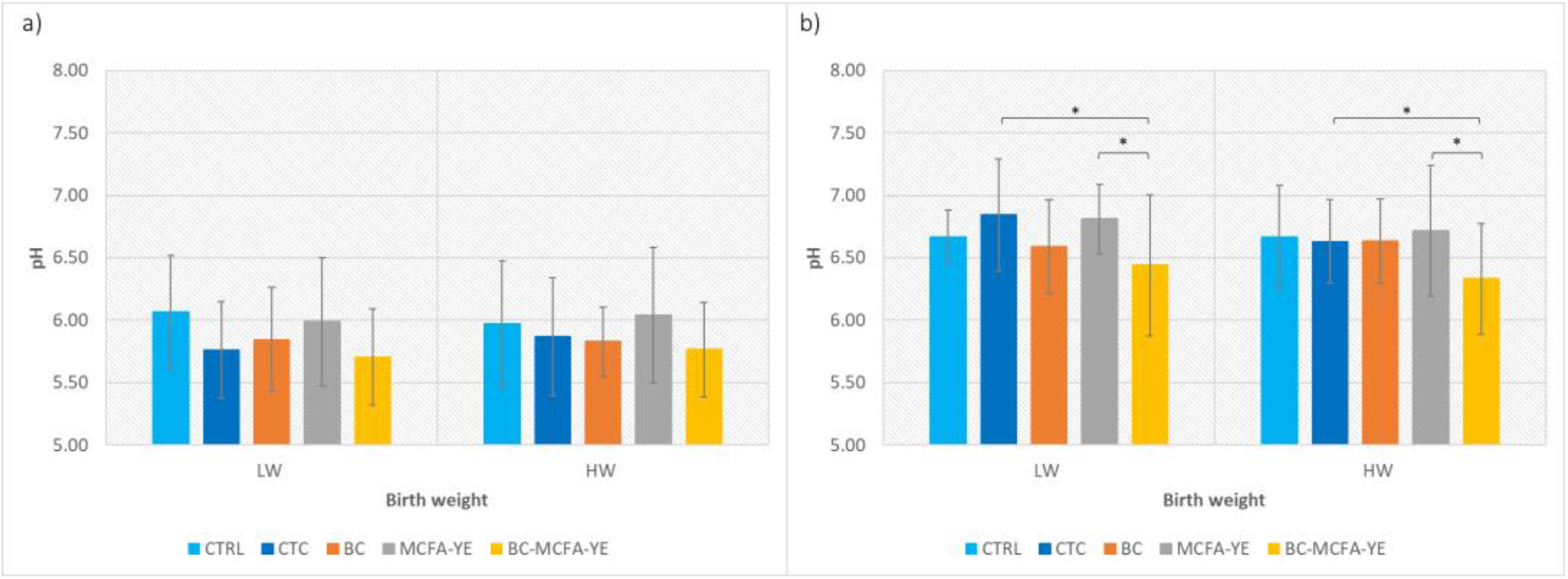
pH of the intestinal content of the (a) caecum and (b) colon of piglets of low birth (LW) and high birth weight (HW) for different feed treatments.

### Volatile fatty acids

The concentrations of acetate, propionate, or butyrate in the cecum and of acetate and propionate in the colon were similar across all piglets (Fig. 3). However, the piglets fed MCFA-YE had lower colon concentration of butyrate than the CTC (P = 0.0202) and BC-MCFA-YE (P=0.0493) piglets (Figure 3f). Birth weight affected the concentration of propionate (P = 0.0323) and butyrate (P = 0.0268) in the cecum and of propionate (P = 0.0179) in the colon.

**Figure 3.**
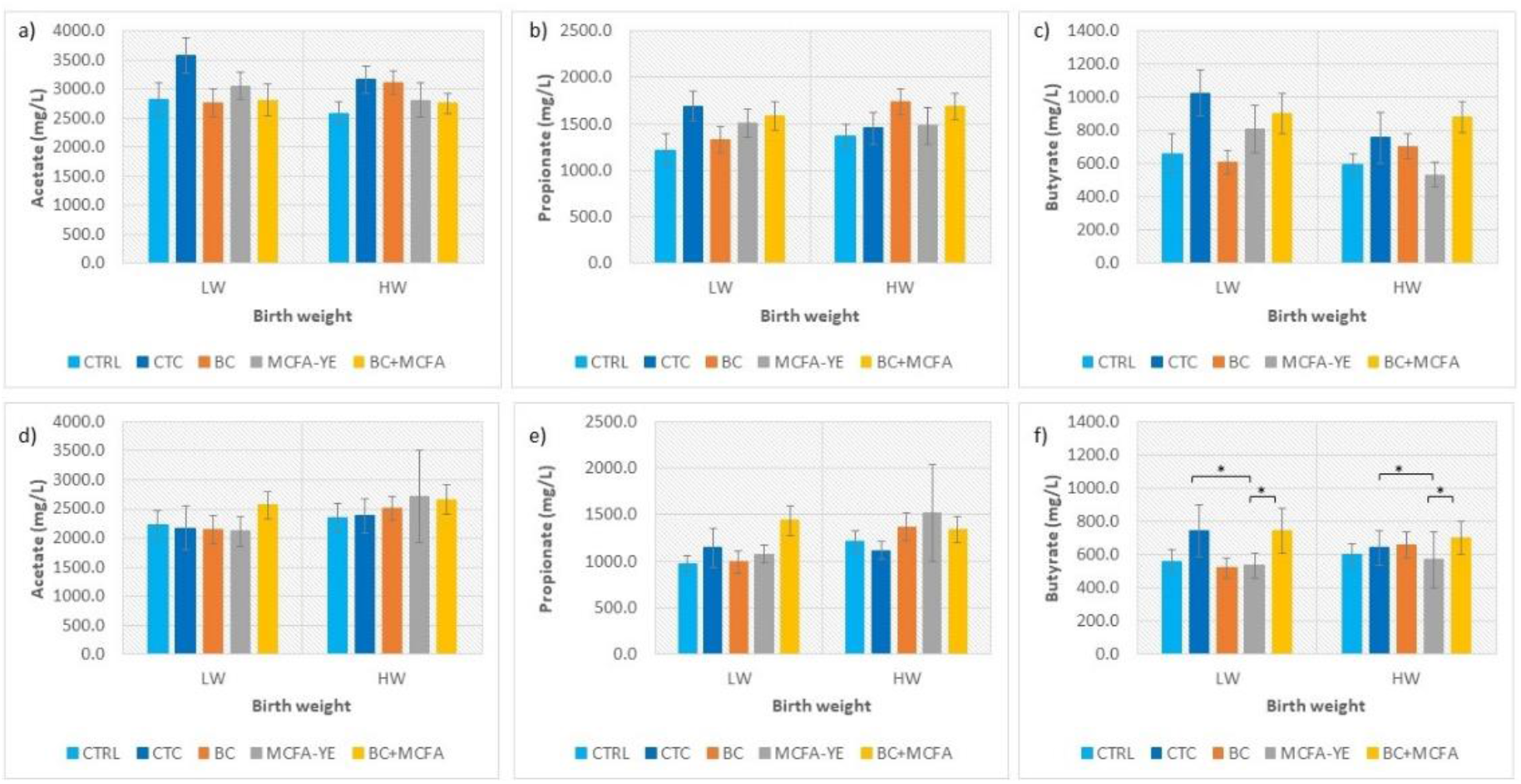
Concentrations of acetate (a and d), propionate (b and e), and butyrate (c and f) found in cecum (a, b, and c) and colon (d, e, and f) digestates of piglet of low birth weight (LW) and high birth weight (HW) for different feed treatments.

### Microbiota diversity, structure and composition

#### Bacterial alpha diversity

The different treatments and the birth weights did not significantly affect the Shannon diversity index nor the number of observed amplicon sequence variants (ASVs), both for the ileum and the cecum (Fig. 4). For both the Shannon index and the number of observed ASVs, the numbers were lower in the ileum as compared to the cecum (Fig. 4).

**Figure 4.**
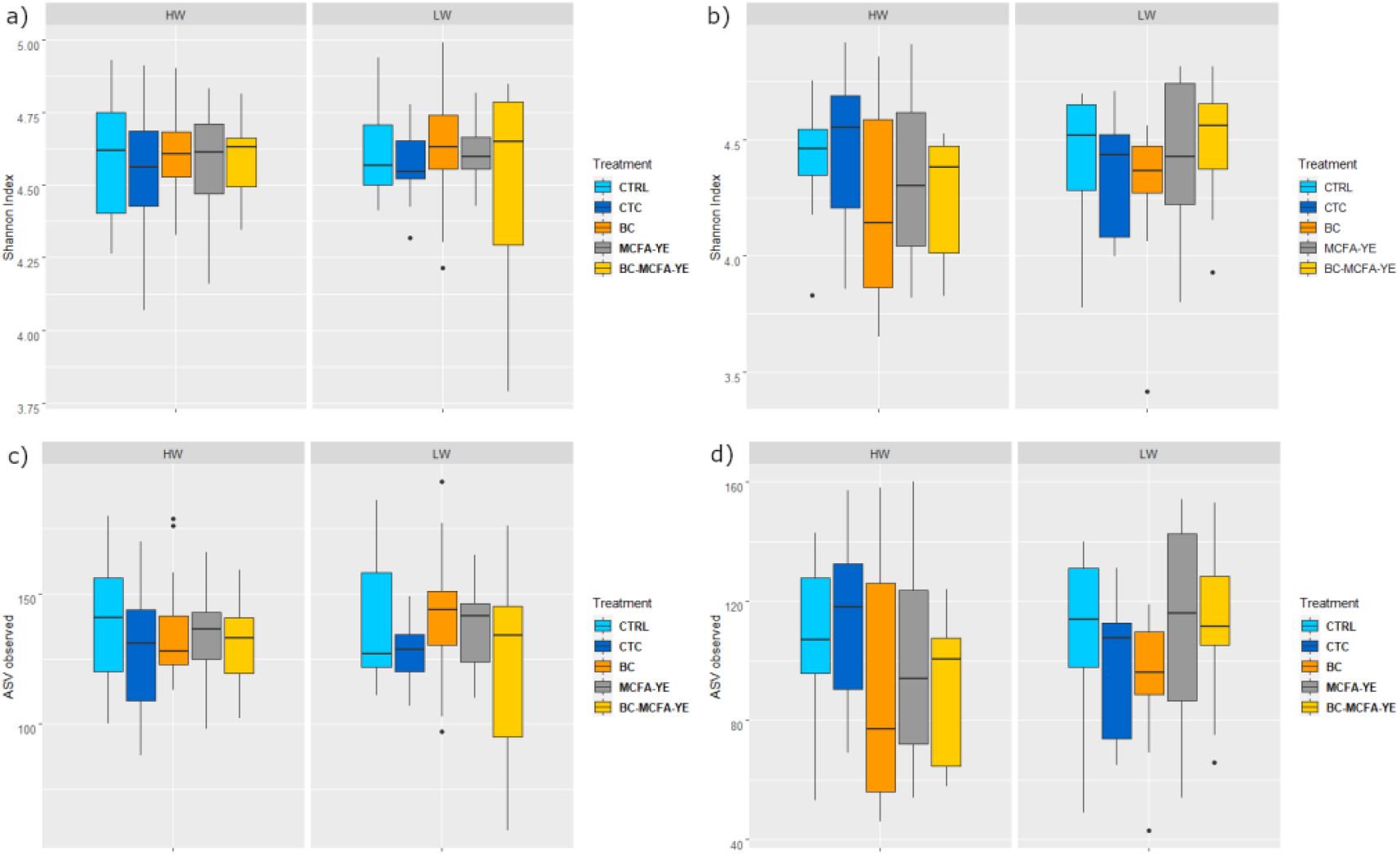
Shannon diversity (a and b) and number of observed ASVs (c and d) based on 16S rRNA gene amplicon sequencing of cecum (a and c) and ileum (b and d) digestate samples from piglets of low birth weight (LW) and high birth weight (HW).

#### Bacterial community structure

Non-metric multidimensional scaling ordinations based on Bray-Curtis’s dissimilarity showed no clear clustering of the samples by treatment or by birth weight (Fig. 5). However, an ANOSIM showed that treatments had significant effects on the bacterial community structure in the cecum (R = 0.0598, P = 0.0005) (Fig. 5a), but not in the ileum (R = 0.00737, P = 0.279) (Fig. 5b).

**Figure 5.**
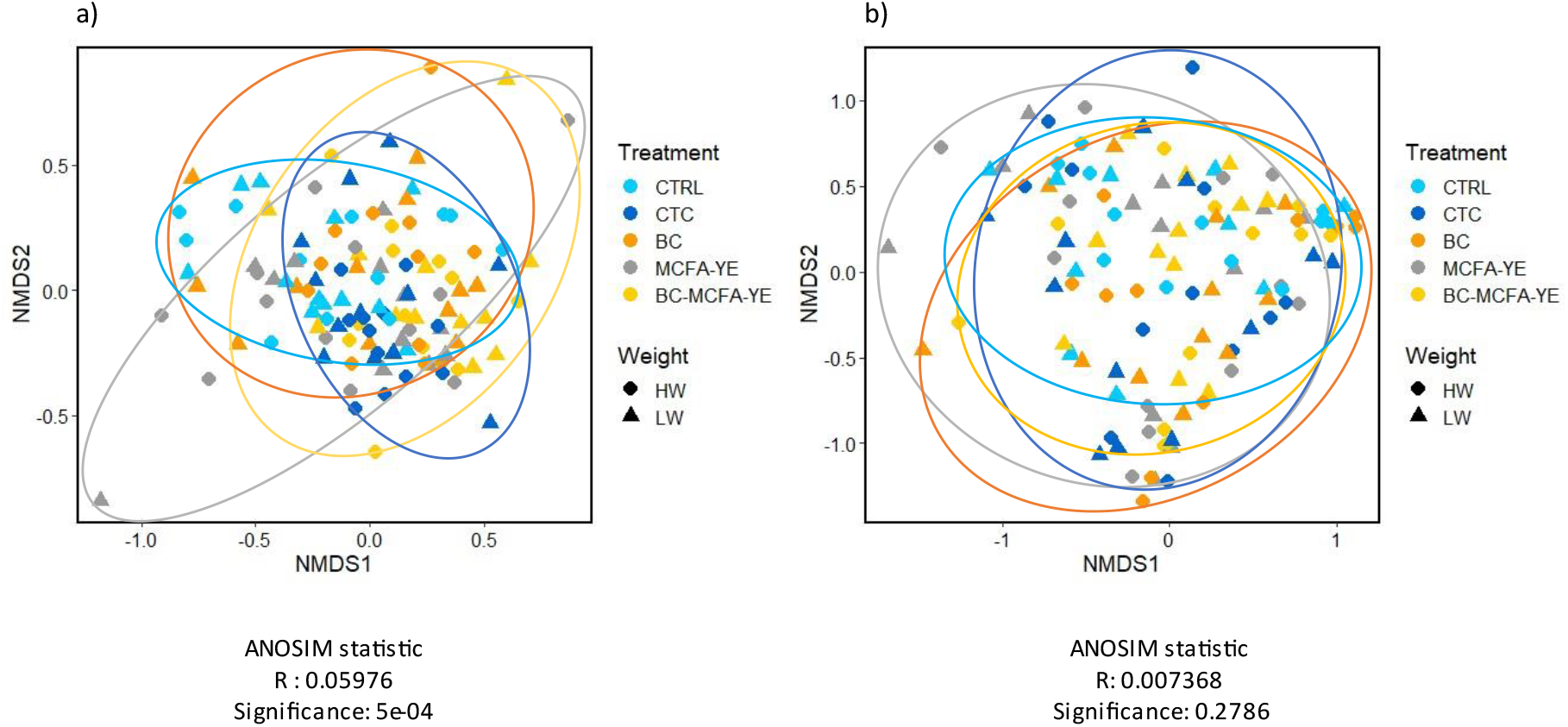
Non-metrical multidimensional scaling analysis (NMDS) and ANOSIM tests based on Bray-Curtis’s dissimilarity at the genus level for cecum (a) and ileum (b) digestates.

#### Bacterial community composition

The relative abundances of abundant (>0.5% of all reads) bacterial phyla and genera present in the cecum and ileum are presented in Figure 6. For the cecum at the phylum level (Fig. 6a), the Bacteroidota and Firmicutes represented more than 95% of the bacterial community, with the Spirochaetota and Proteobacteria phyla also present at a level greater than 0.5%. Pairwise PERMANOVA analyses for the genera relative abundance in the cecum revealed significant differences between the treatments (Fig. 6c). The cecum bacterial genus-level community of the BC-MCFA-YE showed a significant difference from three other treatments, namely the CTRL (P = 0.001), CTC (P = 0.001) and BC (P = 0.019). Furthermore, a significant difference was also observed between the antibiotic controls and the CTRL and BC treatments (P = 0.003 and P = 0.022, respectively). As no significant effect of piglet’s birth weight was recorded, the values were averaged across birth weights.

**Figure 6.**
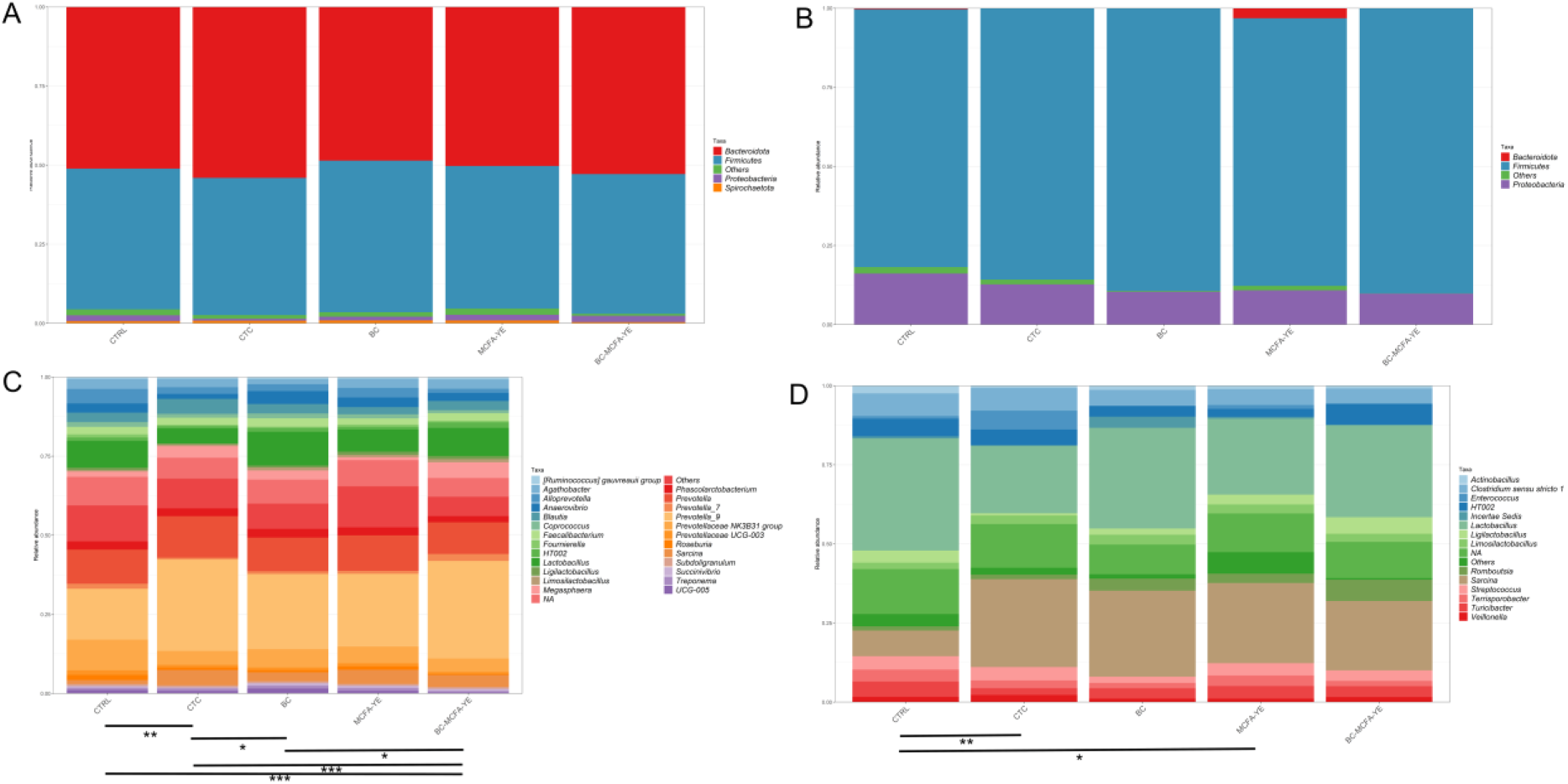
Bacterial community composition at the phylum (a and b) and genus (c and d) levels for cecum (a and c) and ileum (b and d) digestates of piglets subjected to various feed treatments. Values for piglets of low birth weight (LW) et high birth weight (HW) were not statistically different and were averaged for this figure. Only taxa having a relative abundance above 0.5% are shown here. *:P<0.05, **: P<0.01, ***: P<0.001 in pairwise PERMANOVA tests.

As for the bacterial community composition at the phylum level in the ileum, the Firmicutes represented more than 75% of the bacterial phyla detected (Fig. 6b). Proteobacteria represented between 10 to 20% of the bacterial community and Bacteroidota were only present above the 0.5% threshold for the MCFA-YE treatment. At the genus level, pairwise PERMANOVA revealed significant differences between the CTC and CTRL treatments (P = 0.003), and between the CTRL and MCFA-YE treatments (P = 0.049). As for the cecum, there was no significant effect of birth weight on the bacterial community composition in the ileum.

## Discussion

There is a dire need to find alternatives to antibiotics as food supplements in the swine production industry. Here, we compared the effects of three promising supplements, bovine colostrum (BC), medium-chain fatty acids (MCFA), and yeast extract (YE) added to a naked oat base, to a positive control group of animal that ingested chlortetracycline (CTC) and an unamended negative control (CTRL). We measured the piglets weight, intestinal pH, volatile fatty acids (VFA) concentrations and analyzed their gut microbiota. The antibiotic control resulted in the largest shift in the microbial community composition. Significant shifts from the negative control were observed at the genus level for the combined treatment in the cecum and for the MCFA-YE treatment in the ileum. These microbial shifts did not result in significant changes in volatile fatty acid concentrations nor pH values in the cecum or colon when compared to the negative control, although there was a trend toward higher concentrations of VFA in the cecum for the antibiotic treatment, especially for the low-birth-weight piglets. Coherent with these even VFAs concentrations, the piglets of all treatments weighed the same at the time of sampling (day 26). However, the chlortetracycline piglets went on to be heavier than all the other piglets by days 35 and 42. This could be because chlortetracycline increases feed intake (Faccin et al., 2020). In addition to this increase in food intake, it is possible that the trends observed in the microbiota and VFAs concentrations of piglets at day 26 resulted in changes in the weight of piglets later. Alternatively, chlortetracycline might have improved the digestion and, consequently, nutrient uptake by the piglets.

The selected dietary supplements were previously shown to improve swine gut health, robustness, and general performance (Wong et al., 2014;Menchetti et al., 2016;Waititu et al., 2016;Jackman et al., 2020;Lauridsen, 2020;Świątkiewicz et al., 2020). Many of these traits were linked to modulations of the gut microbiota (Frese et al., 2015;Poulsen et al., 2017;Song et al., 2018;Lauridsen, 2020), so the first step was to characterize the effect of these supplements on the piglets gut microbiota as compared to antibiotic and untreated feeds. As expected, and as previously reported (Holman and Chénier, 2014;Ma et al., 2021), chlortetracycline supplementation led to significant shifts in the piglets gut microbiota, both in the cecum and in the ileum. In the cecum, this shift was mainly due to an increase in the relative abundance of some *Prevotellaceae* genera and a decrease in the relative abundance of *Lactobacillus. Prevotella* spp. are generally associated with the consumption of plant polysaccharides, elements found in the post-weaning diet (Ivarsson et al., 2014), and their predominance could favor VFAs production (Frese et al., 2015;Li et al., 2018). The relative abundance of the genus *Sarcina* also increased in the antibiotic treatment. As this genus is often associated with gastric ulcers in humans, dog, and horses and with gastric dilatation that can lead to death in cattle, cats, and horses (Al Rasheed and Senseng, 2016), it could be indicative of a certain level of dysbiosis. The combined treatment (BC-MCFA-YE) led to similarly significant shifts when compared to the antibiotic controls, but without the large reduction in *Lactobacillus* observed in the antibiotic treatment. As *Lactobacillus* are recognized for their potential in preventing infection or colonization by pathogens through competition for nutrients and binding sites in epithelium and by the production of antimicrobial factors such as lactic acid and bacteriocins (Su et al., 2008), their relatively higher abundance in the combined treatment as compared to the antibiotic treatment could indicate an intestinal microbiota that would be more resistant to pathogens. However, this trait was not evaluated in the current study that focused on piglet weight gain. The changes in the ileal microbiota were more subtle and, at the genus level, only the antibiotic and MCFA-YE treatments were significantly different from the control. Both treatments showed increased levels of *Sarcina* and decreased levels of *Lactobacillus*. The antibiotic treatment also increased the relative abundance of *Enterococcus* as compared to the control, which could be explained by the enhanced resistance to antibiotics of the members of this genus (Badul et al., 2021). Taken together our results are showing that some of the feeding treatments did significantly modulate the piglets cecal and ileal microbiota in a way similar to the shifts induced by antibiotic supplements. This is particularly interesting since previous studies had often shown an overriding effect of litter (maternal transmission) or environment on the piglet microbiota around the weaning period, with little to no effects of feed supplements (den Besten et al., 2013;Vigors et al., 2016).

The shifts observed in the cecum microbial community were not mirrored in the VFA concentrations, as no significant differences were observed between the treatments. Shifts in microbial communities measured by amplicon sequencing might not directly translate into functional shifts in the gut functions for many reasons. First, because of functional redundancy, taxonomical shifts might go unnoticed at the functional level. Second, not all the taxa that shifted are involved in the production of VFAs. Third, the taxa that shifted, even if they are known to produce VFAs, might not be actively producing them at the moment of sampling. Fourth, the host uptake of VFAs can vary for numerous reasons. Our results are coherent with previous studies that showed very little changes in cecal VFAs concentration following the use of antibiotics (Peng et al., 2019) and various feed supplements (Awati et al., 2006). However, there were significant differences between some of the treatments in terms of butyrate concentration in the colon. As butyrate is the main energy source for the colonic epithelial cells (Wong et al., 2006), this could impact the energy metabolism of the piglets. These differences seem to have resulted in a lower colonic pH for the BC-MCFA-YE treatment, but not for the antibiotic control, potentially indicating a difference in the absorption of the VFAs. Previous studies have shown that there is not necessarily a link between VFAs concentrations, short-chain fatty acid flux to the host and additional energy harvesting (den Besten et al., 2013), which might explain the lack of coherence of some of the results presented here.

The cecum is the main region of VFA production, where the bacterial community ferments non-digestible carbohydrates and produces metabolites such as short-chain fatty acids (Lauridsen, 2020). These metabolites are essential to the host, including for its energy metabolism (Wong et al., 2006). Coherent with the lack of significant effects in the cecal VFAs concentrations, we did not observe significant differences in the average weight of the piglets on the day of sampling (day 26). However, the antibiotic supplemented piglets were significantly heavier at days 35 and 42, which could be due to the trends observed in the cecal VFA at day 26, with higher concentrations in the antibiotic treatment, especially for the low-birth-weight piglets. It could also have been linked to the shifts observed in the microbiota, especially the increase in the VFA-producing *Prevotella*.

Our results indicate that non-antimicrobial feed supplements can modulate the microbiota of piglets within the first four days after weaning. Although the treatments did not have the expected downstream effects on VFAs production and piglets weight gains, our results are promising as they are showed several important effects on the relative abundance of bacterial phyla and genus in the cecum and ileum. In other words, the microbiota of 4-days post-weaning piglets can be modulated. If this effect can be amplified, the treatments could very well result in the expected changes in piglet weight and provide an efficient alternative to antibiotic supplements. These alternatives are urgently needed.

## Acknowledgments

This study was funded by the Swine Cluster 3 program of the Canadian Swine Research and Development Cluster project #1793.

